# The *C. albicans* virulence factor Candidalysin polymerizes in solution to form membrane pores and damage epithelial cells

**DOI:** 10.1101/2021.11.11.468266

**Authors:** Charles M. Russell, Katherine G. Schaefer, Andrew Dixson, Amber L.H. Gray, Robert J. Pyron, Daiane S. Alves, Nicholas Moore, Elizabeth A. Conley, Tommi A. White, Thanh Do, Gavin M. King, Francisco N. Barrera

**Affiliations:** Department of Biochemistry & Cellular and Molecular Biology, University of Tennessee, Knoxville, TN 37996, U. S. A.; Department of Physics and Astronomy, University of Missouri, Columbia, MO 65211, U.S.A.; Department of Chemistry, University of Tennessee, Knoxville, TN 37996, U. S. A.; Department of Biochemistry, University of Missouri, Columbia, MO 65211, U.S.A.; Electron Microscopy Core, University of Missouri, Columbia, MO 65211, U.S.A.

## Abstract

The pathogenic fungus *Candida albicans* causes severe invasive candidiasis. *C. albicans* infection requires the action of the virulence factor Candidalysin (CL), which damages the plasma membrane of the target human cells. However, the molecular mechanism that CL uses to permeabilize membranes is poorly understood. We employed complementary biophysical, modeling, microscopy, and cell biology methods to reveal that CL forms membrane pores using a unique molecular mechanism. Unexpectedly, it was observed that CL readily assembles into linear polymers in solution. The basic structural unit in polymer formation is a CL 8-mer, which is sequentially added into a string configuration. Finally, the linear polymers can close into a loop. Our data indicate that CL loops spontaneously insert into the membrane to become membrane pores. We identified a CL mutation (G4W) that inhibited the formation of polymers in solution and prevented formation of pores in different synthetic lipid membranes systems. Studies in epithelial cells showed that G4W CL failed to activate the danger response signaling pathway, a hallmark of the pathogenic effect of CL. These results indicate that CL polymerization in solution is a necessary step for the damage of cellular membranes. Analysis of thousands of CL pores by atomic force microscopy revealed the co-existence of simple depressions and complex pores decorated with protrusions. Imaging and modeling indicate that the two types of pores are formed by CL molecules assembled into alternate orientations. We propose that this structural rearrangement represents a maturation mechanism that might stabilize pore formation to achieve more robust cellular damage. Taken together, the data show that CL uses a previously unknown mechanism to damage membranes, whereby pre-assembly of CL loops in solution directly leads to formation of membrane pores. Our investigation not only unravels a new paradigm for the formation of membrane pores, but additionally identifies CL polymerization as a novel therapeutic target to treat candidiasis.

## Introduction

Fungal infections are responsible for high morbidity burden around the world (1). *Candida albicans* is one of the most serious fungal threats to human health. This pathogen is part of the commensal flora, but it can cause infections in the skin, mouth, vagina and gut, both in healthy and immunocompromised individuals. Additionally, *C. albicans* causes invasive candidiasis, an infection of the blood, heart, and other organs, which is common in hospitalized patients (2). Invasive candidiasis causes high rates of mortality (~50%), even when patients are treated with antifungal therapy (2).

It has been recently discovered that *C. albicans* causes toxicity by secreting onto the surface of epithelial cells a peptide toxin, named Candidalysin (CL) (3). This virulence factor binds to the plasma membrane of the target human epithelial cells and compromises the permeability barrier, causing uncontrolled Ca^2+^ influx and release of cellular proteins into the extracellular medium (4). The resulting cellular damage activates a signaling cascade that triggers the release of pro-inflammatory mediators (4, 5), which participate in the immune response to *C. albicans* infection. Furthermore, CL plays a second pathogenic role, as it also attacks the membrane of mononuclear phagocytes, which would otherwise fight the fungal infection (6). CL is necessary for infection by *C. albicans* (7). Therefore, pharmacological inhibition of CL activity is a promising avenue to fight infection of this human pathogen. However, the development of drugs to counter the action of CL is hampered by the lack of understanding of how this peptide damages membranes.

Here we describe how the use of biophysical techniques (mass photometry, native mass spectrometry, analytical ultracentrifugation, transmission electron microscopy, atomic force microscopy, oriented circular dichroism and liposome dye release assay), modeling, and cellular assays allowed us to elucidate that CL damages membranes by forming pores that are assembled using a unique molecular mechanism.

## Results

### CL assembles into polymers in solution

When we ran CL on an SDS-PAGE, we observed not only the expected monomeric band running at ~4 kDa, but also a band of slower electrophoretic mobility (Figure 1A). This result suggests that CL self-assembles in solution, and we hypothesized that this event might be related to its ability to damage cellular membranes. We therefore investigated CL oligomerization, first using native ion mobility-mass spectrometry (IM-MS), a sensitive analytical technique that can isolate and characterize transient oligomers based on their mass-to-charge ratio (*m*/*z*), shape, size, and charge. IM-MS analysis of CL revealed several oligomeric species (Figure 1B). The mass spectral peak at *m*/*z* 3311, which corresponds to a nominal oligomer-to-charge ratio (*n/z*) of *1*/*1*, was notable due to its high intensity when compared to the surrounding oligomers. Analysis of the arrival time distributions (ATD) of this mass spectral peak (Figure 1C) revealed two features that correspond to higher-order CL oligomers. We conservatively assigned the longest arrival time feature (salmon) as an 8-mer with *z* = +*8*. This assignment was performed comparing the isotopic spacing and drift time of the features within the ATD with other mass spectral peaks (see Figure 1 – figure supplement 1). Since a species with a high charge will travel faster than one with a lower charge, the fastest arrival time species (green) is proposed to be a large oligomer resulting from the self-assembly of the 8-mer. The ATD data therefore suggest that CL forms an 8-mer that self-assembles. Given that CL adopts helical structure in solution (3), we built an atomistic model of the 8-mer by aligning CL’s sequence into the eight-helix coiled coil formed by CC-Type2-II (PDB ID 6G67) (1). After the resulting structure was energy minimized (Figure 1D), the theoretical collisional cross section of the model, 2500 Å^2^, agreed with the experimental value of 2567 Å^2^.

**Figure 1.**
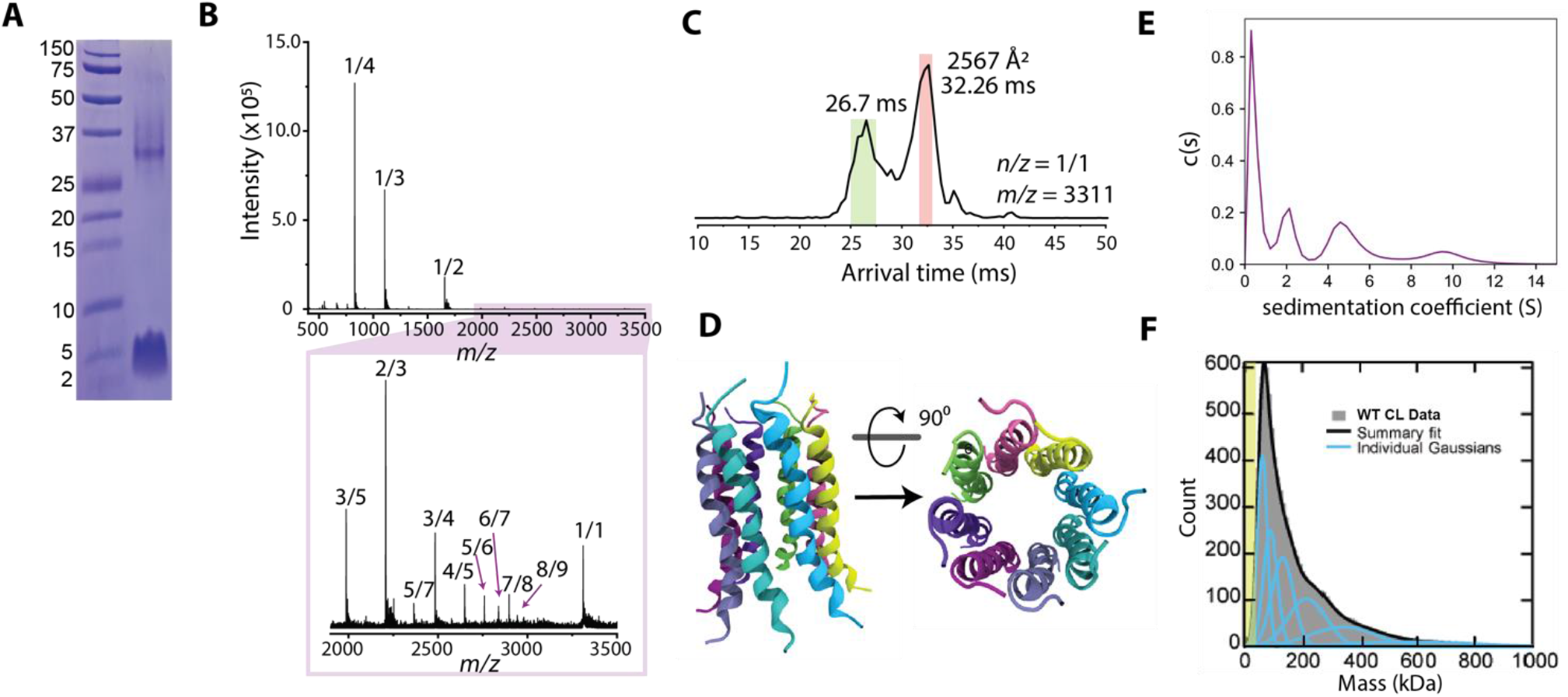
CL forms complex assemblies in solution. (A) SDS-PAGE of CL shows a ~4 kDa monomeric band, and a band corresponding to a large oligomer. Molecular weight markers are shown on the left. (B) IM-MS mass spectrum of CL annotated with oligomer-to-charge (n/z) ratios. The bottom spectrum reveals oligomers that are not immediately identifiable in the top spectrum. (C) Arrival time distribution of the 8-mer (n/z= 1/1) species. This peak was more populated than those corresponding to smaller oligomers, which might result from decomposition of the 8-mer in the gas phase. The experimental collisional cross section for the single 8-mer (highlighted in salmon) is given. (D) Atomistic model of the CL 8-mer, with a molecular weight of 26.5 kDa. Each monomer helix is colored differently. (E) Analytical ultracentrifugation data identifying populations of CL oligomers of increasing sizes. (F) Mass photometry data of CL oligomeric species. The green area marks the approximate mass range (<50 kDa) below the resolution of the technique. Data distribution was best fit with 6 Gaussian populations (shown as blue lines, and summary fit is the black line). The mass of the peaks agrees with the expected mass for two, three, five, eight, thirteen and twenty-two CL 8-mers.

We used two additional biophysical methods to confirm and further investigate CL oligomerization in solution. We first performed analytical ultracentrifugation of CL (Figure 1E). The sedimentation velocity results reveal a low sedimentation peak, likely corresponding to a CL monomer (8), and several larger assemblies, in agreement with the IM-MS data analysis. We next used mass photometry (MP), a single-molecule technique that is suited for the study of large protein assemblies. While MP is insensitive to the mass of the CL 8-mer (26.5 kDa) as the resolution of the instrument is limited to particles >50 kDa, MP has high sensitivity to larger species. Figure 1F displays MP data of CL, consisting of a main peak and a long tail. The main peak corresponds with the mass of two bound 8-mers, while the larger mass peaks correspond to the progressive assembly of 8-mers (see Figure 1 – figure supplement 2). The long tail of the MP data reaches beyond 600 kDa, which would correspond to the assembly of tens of 8-mers. Taken together, the four techniques reveal that CL readily forms large oligomers in solution and suggest that the 8-mer is the seed for further CL self-assembly into large structures.

We employed microscopy to resolve the assemblies that CL forms in solution. We first performed negative-stain transmission electron microscopy (TEM), and observed that CL does not form amorphous aggregates, but instead it assembles into linear structures (Figure 2A). The TEM data revealed a basic structural unit, which seemed to grow in a step-wise fashion into polymers (Figure 2A, side panels). The longer polymers curved, and in some cases closed in on themselves, forming a loop with diameter ≥10 nm. We further studied CL polymerization by atomic force microscopy (AFM), since this technique provides high resolution images of single particles and can also be readily used to image membrane pores. AFM images of CL in buffer (Figure 2B) agreed with the TEM results and confirmed that CL polymerizes and can form loops. The bending stiffness of CL polymers was calculated from AFM images through persistence length, *L_p_*, analysis (*L_p_*= 9 nm ± 2 nm, *N* = 100, Figure 2 – figure supplement 1). The bending stiffness of linear CL polymers is significantly lower than other biological polymers, like actin fibers (9), explaining the ability of CL polymers to close into loops.

**Figure 2.**
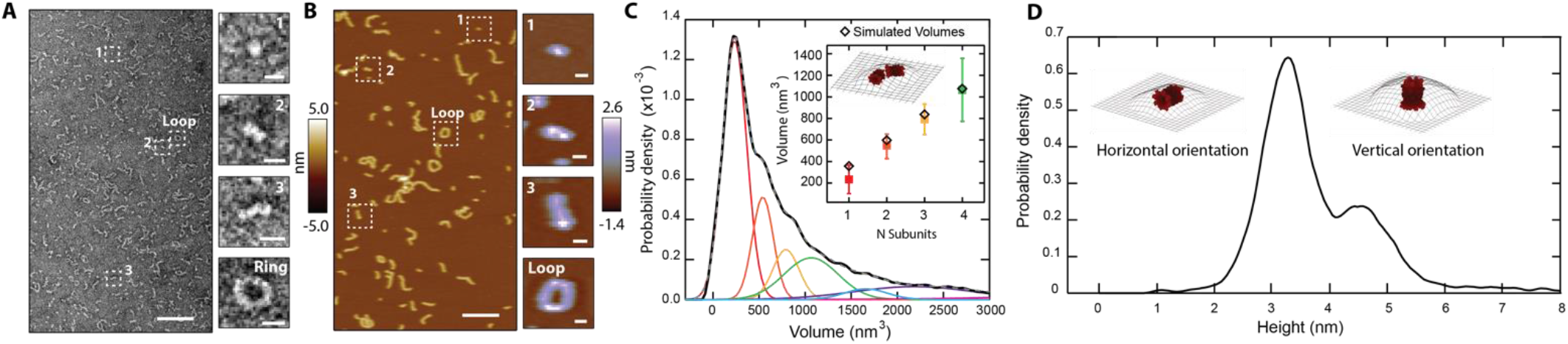
CL in solution forms progressively longer linear polymers and loops. (A) TEM reveals CL polymerization; scale bar = 100 nm. Features of growing complexity are magnified in the side panels; scale bar = 10 nm. (B) AFM imaging in fluid shows overall agreement with TEM data; scale bar = 100 nm. Four similar CL features are also highlighted in the side panels; scale bar = 10 nm. (C) Smoothed volume histogram of *N* = 7,838 individual features (solid black line) fitted with Gaussian distributions (solid colored lines, summary fit is the gray dashed line). The inset compares the experimental volume of the first four peaks (squares colored as in panel C) with simulated volumes (black diamonds) calculated from *N* 8-mers. The cartoon shows an example of head-to-toe assembly of two 8-mers. Error bars for the peak positions represent the standard deviation. (D) When the height of the individual features was measured, it yielded a bimodal distribution, with heights of 3.2 nm and 4.6 nm. The insets show the two proposed orientations of the 8-mer. The topographic surface is overlaid to show convolution of the AFM tip.

Examination of AFM images containing thousands of particles yielded a volume distribution with a sharp peak and a long tail (Figure 2C), which resembles the distribution observed in the MP data (Figure 1F). Such agreement suggests that the AFM results are robust and are not caused by spurious interactions with the AFM tip. The AFM volume histogram was deconvolved by fitting Gaussian distributions, revealing a primary peak and several larger sub-populations of larger volumes (Figure 2C). The volume increased in approximately constant steps between the populations (see side panels of Figure 2B), confirming that a basic structural unit grew first by dimerization, followed by sequential addition to further increase linear polymer length.

We measured the height of the individual particles observed by AFM, and the resulting histogram (Figure 2D) yielded two peaks, with heights of 3.2 ± 0.4 nm and 4.6 ± 0.5 nm. We reasoned that the bimodal height distribution could indicate two orientations (vertical and horizontal) of the basic structural unit (Figure 2B inset, labeled “1”), which is probably the CL 8-mer identified by IM-MS (Figure 1B). We tested this possibility by using the CL 8-mer model (Figure 1D) to generate simulated AFM images through morphological dilation of the estimated tip geometry for the 8-mer, either on its horizontal or vertical orientations (Figure 2D, insets). We found good agreement between the simulation [*heights* of 3.3 nm and 4.8 nm] and the two experimental peaks shown in Figure 2D. Interestingly, the *volume* of the modeled 8-mer in the horizontal orientation additionally agrees within uncertainty with the value of the basic AFM structural unit (Figure 2C inset). Taken together, these results support the notion that CL assembles into an 8-mer in solution.

Polymer growth of CL 8-mers in the horizontal orientation (lying flat on a surface) can occur by addition of a second 8-mer in two fundamental modes, side-by-side or head-to-toe. When we modeled the latter arrangement for the assembly of two, three, and four head-to-toe 8-mers (Figure 2C, inset), we found agreement between the simulated and experimental AFM volumes (Figure 2C, inset; Figure 2 – figure supplement 2). Taken together, the data indicate that a CL polymer is formed when a basic subunit, which our results indicate is an 8-mer, is aligned in the longest dimension, and grows by sequential head-to-toe addition of additional 8-mers.

### CL forms two classes of membrane pores

CL mediates *C. albicans* infection by damaging the integrity of the plasma membrane of human cells (3). It has been proposed that CL forms membrane pores (3), but such structures have never been observed. We performed AFM imaging on supported lipid bilayers made of DOPC to gain insights into how CL causes membrane disruption. We observed that in the presence of CL, bilayers indeed exhibited punctate depressions commonly associated with pores (Figure 3A). Though the AFM tip is sharp (nominal radius ~8 nm), the pores’ radii were often smaller that the tip, preventing passage all the way through the 4-nm-thick membrane, which cause artificially shallow readings (10, 11) (Figure 3-figure supplement 1).

**Figure 3.**
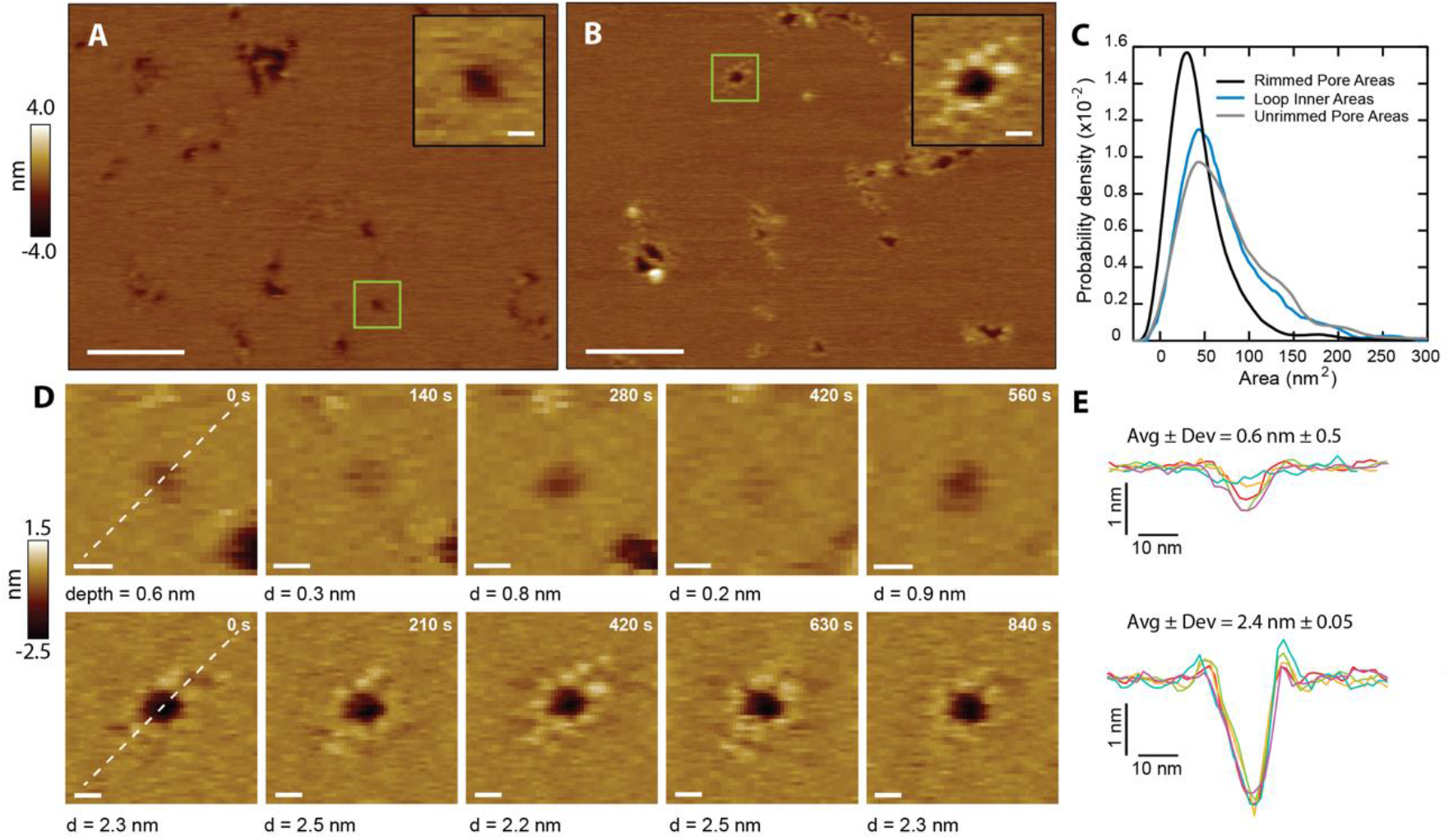
CL forms two types of membrane pores. Representative images showing predominantly unrimmed (A) and rimmed (B) pore features in supported DOPC membranes; scale bars = 100 nm. The insets show zoomed views of green boxed features; scale bars = 10 nm. (C) Area histograms show that unrimmed pores (gray line) display a broad peak at 41 ± 25 nm^2^ (mean ± S.D.) with a shoulder at ~130 nm, similar to loops found in solution (38 ± 21 nm^2^, aqua blue line). In contrast, the rimmed pores (black line) exhibit a narrower distribution with a smaller area (26 ± 19 nm^2^) (*N_unrimmed pores_* = 1,468, *N_rimmed pores_* = 492, *N_loops_* = 261). (D) Unrimmed (*top*) and rimmed (*bottom*) features were imaged over several minutes; scale bars = 10 nm. (E) Line scans, marked as white lines in the previous panel, show the dynamics of the pore profile in both cases. Different colors were used for each image. The average depth (Avg) and relative deviation (Dev), defined as the standard deviation of the depth divided by Avg, are listed.

The CL pores fell into two broad categories: simple depressions (Figure 3A), and complex pores surrounded by a rim of discrete protrusions of a height of 0.3 nm above the bilayer surface (Figure 3B). To differentiate between the two types of pores, they are henceforth referred to as “unrimmed” (Figure 3A) and “rimmed” (Figure 3B) pores, respectively. Both types of pores were free to diffuse in the bilayer (Figure 3-figure supplement 2), indicating that CL is not immobilized by interactions with the mica surface. Rimmed pores generally have a smaller area than unrimmed pores (Figure 3C). Unrimmed pores have a broad area distribution, with the main population at 41 nm^2^ (Figure 3C, gray), while the rimmed pores have a narrower distribution, and an area peak at 26 nm^2^ (Figure 3C, black). Selected areas of the samples were imaged repeatedly to produce a time series of both unrimmed and rimmed pores. A representative unrimmed pore (Figure 3D, top row) shows dynamic behavior, with large variations over time. The pore depth visibly varies, flickering between deep and shallow states. On the other hand, the rimmed pore (Figure 3D, bottom row) appears deeper, and is more stable, as shown by the constant profile over time (Figure 3E). These results suggest that the presence of the rim endows the pore with stability, which could potentially be a mechanism to afford CL pores with enhanced membrane damaging capabilities.

### CL polymers insert into the membrane, and loops become membrane pores

We noticed that there were strong similarities between the shape and size of the loops that CL forms in solution and the unrimmed membrane pores (Figure 3C, compare blue and gray lines). This agreement led us to hypothesize that CL loops could insert into membranes and become unrimmed pores. We could test this hypothesis as occasionally a patch of supported bilayer would spontaneously dissociate from the underlying mica surface (Figure 4A). Upon imaging the exact same area again, we observed features similar to the CL polymers and loops detected in solution. Indeed, loops, often inter-connected, were observed where pores were previously present (see blue arrows). Linear features similar to the CL polymers observed in solution were also present, which had been previously obscured by the presence of a bilayer. This observation indicates that linear polymers can insert into membranes without causing membrane disruption, but when they close into a loop they form an unrimmed membrane pore.

**Figure 4.**
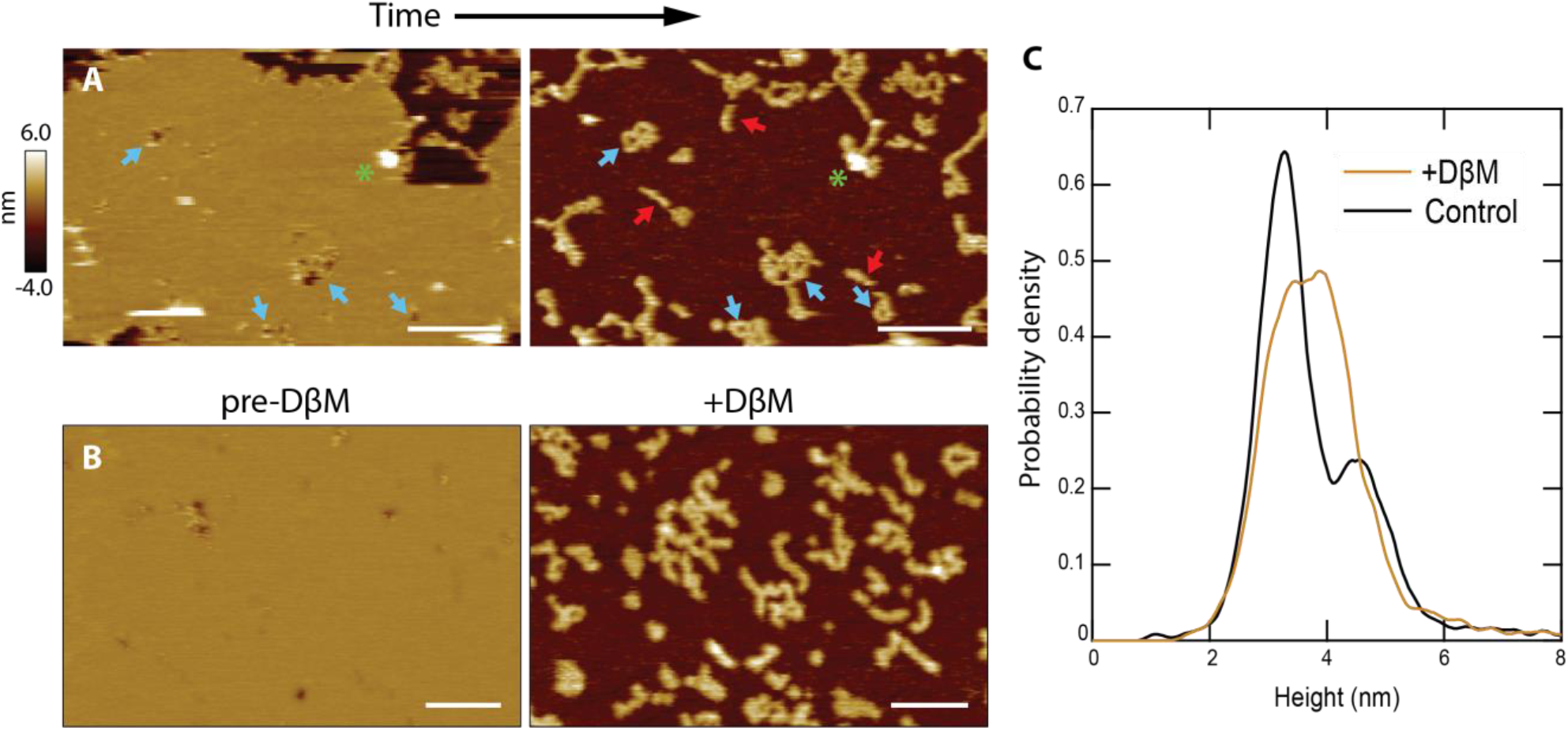
CL polymers and loops insert into membranes. (A) Lipid patches exhibiting pore-like features occasionally dissociated from the mica surface, revealing underlying structures similar to CL polymers in solution [see Fig. 2B]; scale bars = 200 nm. Blue arrows indicate pore features and corresponding polymer loops/tangles. Linear features were revealed that were not previously observed when lipid was present (red arrows). A green asterisk draws attention to a tall positive feature that remains in both images, serving as a reference. (B) Detergent was used to forcibly remove lipid bilayers. The addition of DβM followed by rinsing revealed underlying CL structures. Scale bars = 100 nm. (C) A histogram of particle heights compares the solution data (black, *N* = 7,838) to the features remaining after addition of DβM to the CL+DOPC system (gold, *N* = 8,170). The solution results were bimodal, indicating two overall 8-mer orientations. The leftover features after detergent removal of the membrane exhibited a broad peak roughly encapsulating both of the solution peaks.

In order to replicate these spontaneous events in a deterministic manner, we used the mild detergent dodecyl beta-D-maltoside (DβM) to remove the lipid bilayer (12, 13). Once the detergent and solubilized lipid were rinsed away, the CL structures remaining were imaged (Figure 4B). Control experiments were performed that showed that DβM efficiently removed DOPC molecules from the mica support, but did not dissociate the CL polymers (Figure 4-figure supplement 1). The histogram in Figure 4C reveals similarities between the height of CL features in solution, showing a bimodal population, and in membrane areas post-DβM treatment. In these latter samples there was a broadening of the primary height peak, which can be attributed to binding between CL and the detergent. However, the increase in taller features could also be an indication that the vertical 8-mer orientation (**Fig. 2D**) is favored in the presence of lipid membranes.

Our data indicate that loops can insert into membranes and form unrimmed pores. These might later mature into the rimmed pores, which have better defined dimensions. The question then arises regarding what orientation, head-to-toe or side-by-side, does the CL 8-mer adopt in the two types of pores. To investigate this question, we modeled loops of *N* subunits in the two 8-mer orientations, head-to-toe and side-by-side (Figure 5A). Comparison between the experimental areas and the models (Figure 5A, Figure 5, figure supplement 1) suggest that the *loops* found in solution, similarly to the *linear polymers*, assemble into a head-to-toe fashion. The average loop would be composed of six head-to-toe 8-mers, for a mass (159 kDa) that corresponds with the start of the tail in the MP distribution (Figure 1F).

**Figure 5.**
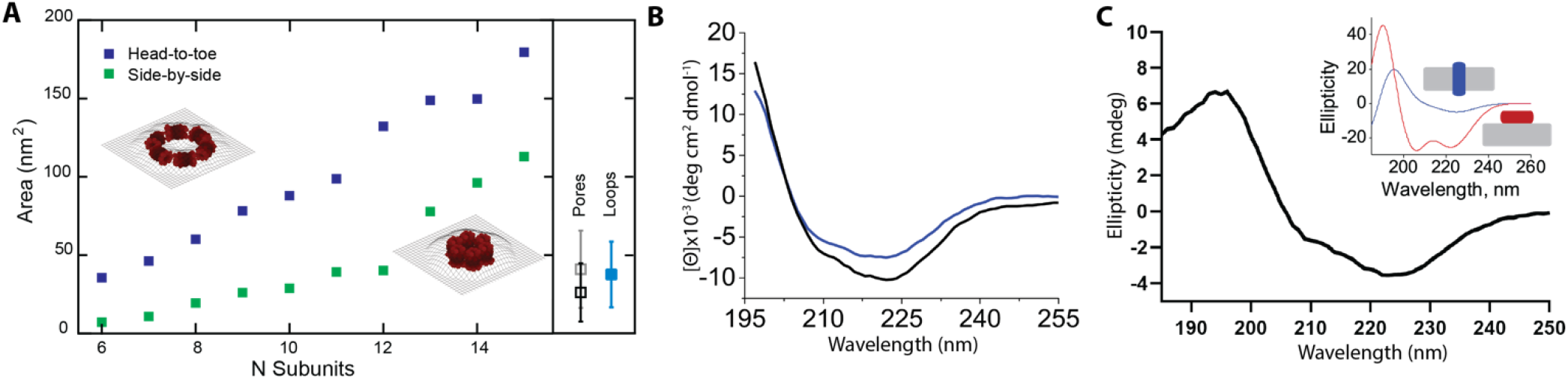
Rimmed pores appear upon a conformational rearrangement. (A) CL pores were simulated using two possible arrangements of the 8-mer subunit, side-by-side (green) and head-to-toe (dark blue) 8-mer. The particle area is plotted versus the number of subunits. The insets show simulations for N=6 in both orientations. Closed loops with less subunits were not geometrically likely. The side panel shows the experimental areas for unrimmed pores (gray symbol), rimmed pores (black), and the loops found in solution (aqua blue); error bars represent standard deviation. (B) Standard circular dichroism (CD) of CL in buffer (blue) and in the presence of lipid vesicles (black). (C) OCD data in supported membranes indicate that CL adopts largely a TM orientation, but it also suggests the presence of a population of CL molecules with helices aligned with the membrane plane. The inset shows theoretical OCD curves for TM helices (blue) or surface helices (red).

We determined the areas of the membrane pores taking advantage of the features that remained after spontaneous lipid dissociation (Figure 4A). For unrimmed pores, the AFM data also corresponded to a head-to-toe 8-mer arrangement (Figure 5A). Loops with a side-by-side orientation were rare in the solution data, but were common in CL samples that had been exposed to lipid. Indeed, comparison between the model and the rimmed pores data suggest that in these pores the 8-mers switch from the head-to-toe orientation found in the unrimmed pores (gray square, side panel of Figure 5A), to a side-by-side orientation (black square).

Specifically, the modeled areas predict that unrimmed pores contain six to eight head-to-toe 8-mers, while rimmed pores are formed by six to twelve side-to-side 8-mers. Therefore, we propose that a rimmed pore (Figure 3B) appears when the 8-mer subunits in an unrimmed pore rotate 90 degrees, switching from the head-to-toe to the side-by-side arrangement (compare insets in Figure 5A).

We interrogated the results of the modeling using circular dichroism (CD). Figure 5B shows that CL forms, as expected (3), an α-helical structure in solution, as evidenced by the spectral minima at 208 and 222 nm, and that the presence of lipid vesicles promote modest additional helix formation. The orientation of the CL α-helix with respect to the membrane plane can be determined using oriented CD (OCD), which is performed using supported membranes (14). OCD can discriminate between helices aligned along the membrane plane and transmembrane (TM) helices (15) (Figure 5C, inset). The OCD data of CL displayed low intensity, and a minimum at ~225 nm and a maximum at ~195 nm, similarly to the expected result for TM helices (16, 17). The OCD spectrum therefore indicates that the predominant orientation of helices in CL in membranes was inserted across the membrane, in a TM orientation. This is the proposed helical alignment that is expected in the side-by-side 8-mer arrangement of the rimmed pores. However, the presence of a depression at ~210 nm agrees with the presence of a population of α-helices oriented along to the plane of the membrane, like those expected for the unrimmed pore. Therefore, the OCD result supports the presence of a combination of rimmed and unrimmed CL pores with the respective helical orientations predicted by the AFM modeling.

### Mutational analysis of CL provides mechanistic insights

Our experimental data and modeling indicate that CL 8-mers in solution polymerize in a head-to-toe fashion. We sought to further validate this hypothesis by performing mutations in the CL sequence. A head-to-toe assembly of a parallel helical bundle implies interaction between the N-terminal (N_t_) helical ends of an 8-mer with the C-termini of the adjacent subunit (Figure 6-figure supplement 1). Our 8-mer model predicts that the Nt is slightly kinked (Figure 1D), and glycine residues confer flexibility to α-helices (18). We reasoned that mutation of residue G4 (Figure 6A), located at the Nt kink, might hamper CL polymerization. We replaced this residue with a bulky tryptophan side chain to maximize the expected disturbance. We first tested the ability of the resulting G4W CL variant to polymerize in solution using MP, and observed that G4W formed few large assemblies (Figure 6B). The residue G4 lies at the outer surface of the hollow cylinder that the 8-mer forms. We sought to test the effect of a residue at the opposite side of the helix, which is predicted to form the core of the structure. We therefore tested the I9A mutation. The MP results show that the I9A variant, on the other hand, self-assembles efficiently in solution (Figure 6B).

**Figure 6.**
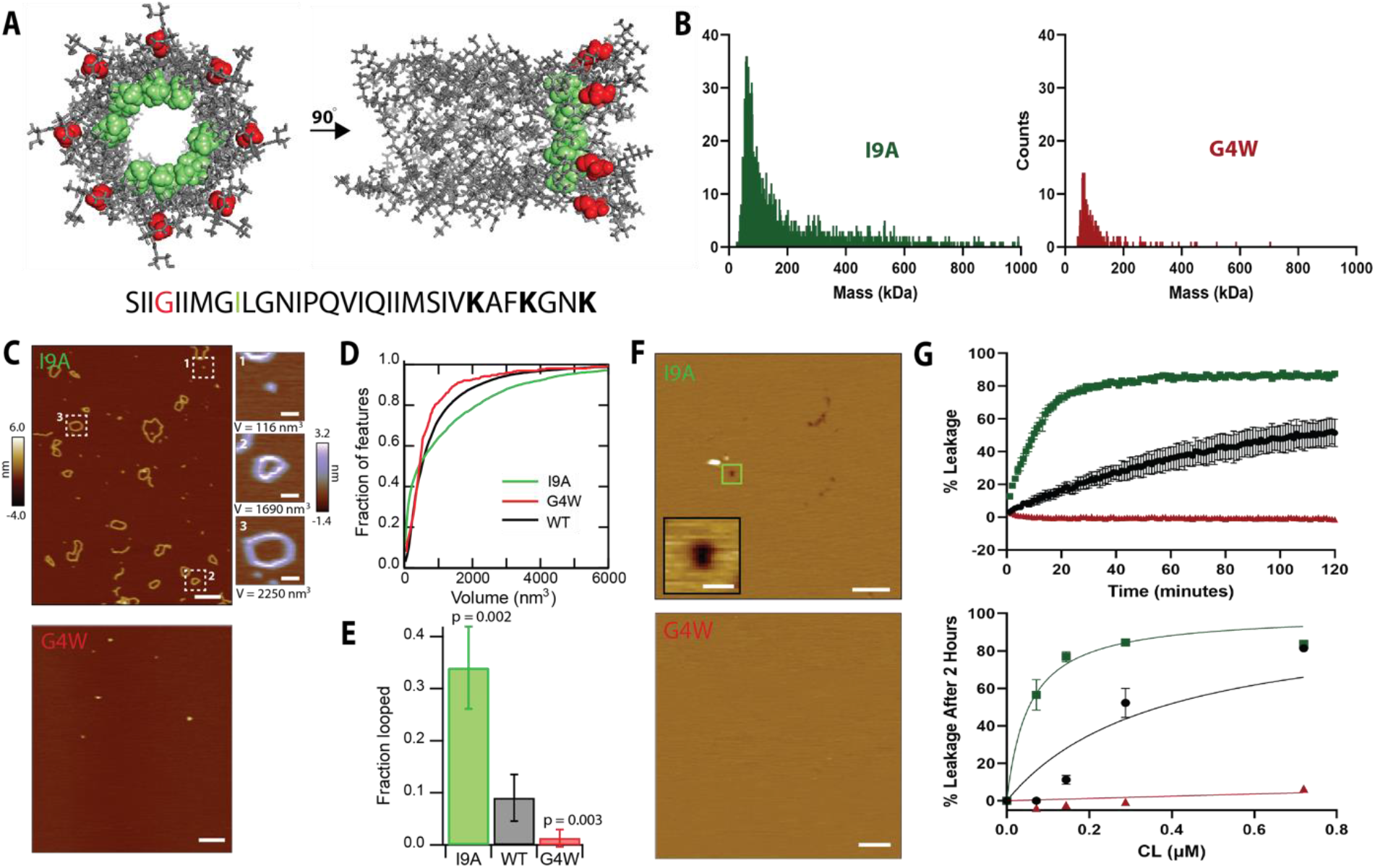
Mutational analysis indicates that polymerization in solution is required for pore formation. (A) The position of the residues G4 (red) and I9 (green) are shown for two different 8-mer orientations. The CL sequence highlighting the position of the mutations is shown at the bottom. (B) Mass photometry shows that the G4W variant has a lower tendency to form large assemblies than I9A in solution. (C) AFM data of CL variants in solution. The I9A variant exhibits increased loop features, while G4W does not form polymers; scale bar = 100 nm. Representative I9A features are selected from the image: a protomer, a loop, and a large loop; scale bars = 10 nm. (D) The volumes of the features observed by AFM were calculated, and are shown as accumulated fraction. Data are shown for CL WT (N = 7,838 features), I9A (N = 3,609), and G4W (N = 190). (E) The fraction of all polymers that close into a loop was calculated for all variants. Error bars are standard deviations (*n*= 3 independent experiments); p-values are the result of a Student’s t-test comparing the I9A and G4W data to the WT. (F) I9A CL forms membrane pores, while pores were not observed for G4W; scale bars = 100 nm for full images, 10 nm for inset. (G) Liposome dye release assay shows that CL variants display different membrane disruption. *Top* panel shows a time course of dye release for CL WT (black), I9A (green), and G4W (red) at ~0.3 μM, and the *bottom* panel shows the percentage of dye release after two hours for different peptide concentrations. N = 3-4 and error bars are S.D.

We next tested the two variants using AFM to visualize the morphology of the polymers they formed. The AFM results agreed with the MP, as I9A formed long polymers and closed loops, while G4W formed few large structures (Figure 6C, compare top and bottom panels). We quantified the AFM images by measuring the volume of all particles, and compared these results with AFM data of WT CL. The resulting Figure 6D shows that G4W does not efficiently assemble into large (> 1,000 nm^3^) structures, while WT and I9A do. However, only the I9A variant forms a significant number of very large assemblies (>3,000 nm^3^), which generally corresponded with over-sized loops (Figure 6C, side panel). Since the AFM data (Figure 4) indicate that linear polymers need to close to form membrane pores, a key parameter for membrane disruption would be the ability of polymers to close into loops. We determined the frequency of loop formation for the three peptides (Figure 6E), and observed that there is overall agreement between the ability to form polymers and the loop formation propensity. Consistent with the model that loops in solution become pores, we observed that I9A formed larger membrane pores than WT (Figure 6-figure supplement 2), while we did not observe membrane pores formed by G4W (Figure 6F). Additionally, the higher I9A pore area has a corresponding increase in the inner area of the loops compared to WT CL (Figure 6– figure supplement 2), supporting the hypothesis that the pore architecture is defined by the loops.

We studied next the ability of CL WT and the two variants to disrupt the integrity of POPC vesicles in suspension. Specifically, we carried out an assay where membrane damage is measured by the fluorescence de-quenching that occurs when the dye calcein is released from vesicles. Figure 6G shows the kinetics of membrane disruption caused by the addition of WT and the two CL variants. We observed that G4W did not induce membrane disruption, in agreement with the lack of membrane pores observed by AFM. CL WT efficiently disrupted membranes, but I9A was the most efficacious at disrupting membrane integrity, as expected from the high loop formation probability (Figure 6E) and the presence of larger membrane pores (Figure 6– figure supplement 2). The differences in leakage were consistent over a variety of peptide concentrations in the nanomolar range (Figure 6G). Overall, the mutational data confirm that CL uses a novel molecular mechanism, where it first self-assembles in oligomers in solution, which in turn form polymers that close forming a pore-competent soluble loop (Figure 8). The results with the loss-of-function G4W variant revealed that formation of polymers in solution is a necessary step for the formation of membrane pores.

CL pores damage the plasma membrane of epithelial cells. This attack triggers the activation of danger response signaling, which induces phosphorylation of MAPK phosphatase 1 (MKP1) and subsequent overexpression of the transcription factor c-Fos (5, 19). When we treated the oral epithelial cell line TR146 with WT CL, we observed robust increases in c-Fos (3-fold) levels and phosphorylation of MKP1 (5-fold) and (Figure 7), as expected (3). The I9A variant also induced strong MKP1 phosphorylation, but significantly higher c-Fos expression. Conversely, G4W CL failed to activate the danger response signaling pathway altogether, revealing an overall agreement between the effect of CL mutations in biophysical and cellular assays. Our data therefore indicate that mutations that alter polymerization impact epithelial cell damage response. Taken together, these results suggest that polymerization of CL in solution is a major factor defining *C. albicans* infection.

**Figure 7.**
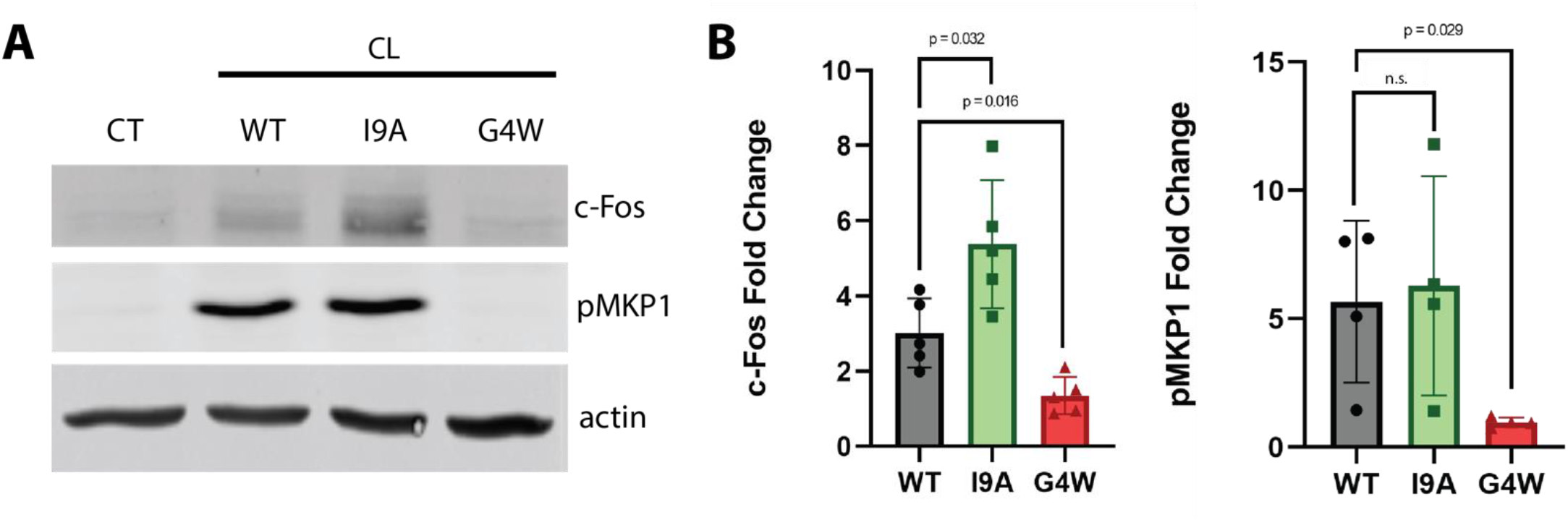
The polymerization-defective G4W variant does not cause danger response signaling in epithelial cells. (A) Representative Western blots of oral epithelial cells (TR146) probed for c-Fos expression and phosphorylation of MKP1 after 2-hour treatment with WT and variant CL. (B) Quantification of c-Fos and MKP1 phosphorylation was used to assess danger response signaling. Data were normalized to values obtained in control conditions. Actin was used as a loading control. N = 4, and bars are S.D.

## Discussion

Pore-forming toxins (PFT) populate different states on their way to form membrane pores. PFT typically are found as soluble monomers that later bind to the lipid bilayer. *After* membrane binding, PFT monomers find each other and oligomerize (20, 21), often forming membrane-anchored pre-pores that later transition into the mature pores that permeabilize membranes (22, 23). Here we show that CL follows a different molecular mechanism for pore formation. Multiple lines of evidence indicate that CL forms both linear and closed loop polymers *before* binding to the membrane (Figure 8). Further, acting as soluble pre-pores, closed loop CL polymers insert into lipid bilayers to cause membrane damage without any observable conformational change.

**Figure 8.**
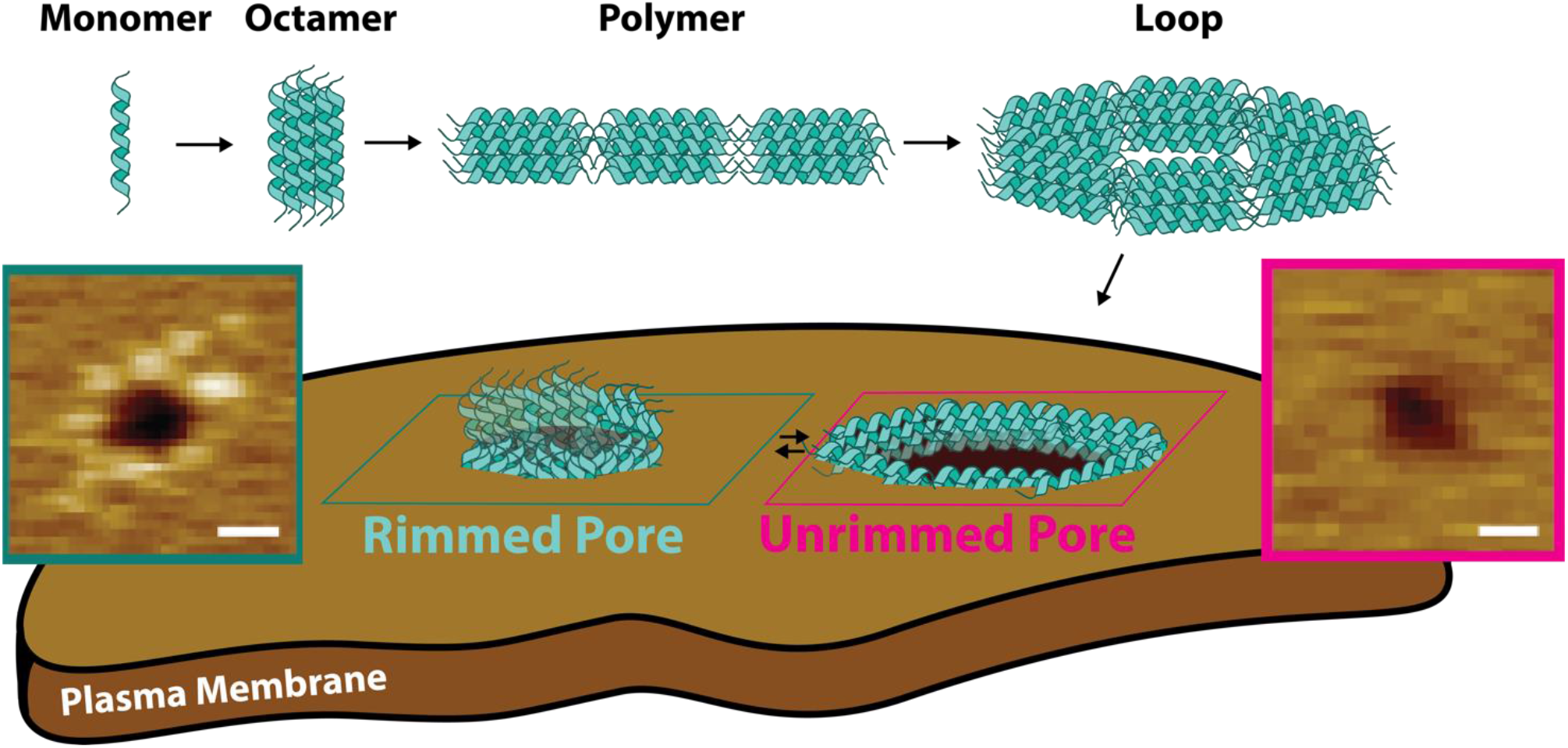
Candidalysin utilizes a novel mechanism of pore formation to attack human cell membranes. The CL monomer oligomerizes into an 8-mer. CL polymers are formed when 8-mers assemble in a head-to-toe fashion. Polymers can close into loops, which have the ability to insert into membranes forming unrimmed pores. Upon insertion, pores exist in a dynamic equilibrium between unrimmed and rimmed pores, which cause cellular toxicity.

It is interesting to compare the topography of CL pores to those formed by other PFTs. AFM imaging revealed that the peptide CL (3.3 kDa) assembles into large membrane pores, of a diameter that is similar to the aqueous pores formed by the proteins gasdermin-D (24), suilysin (20), and perfringolysin O (25, 26), of molecular weights in the 50-60 kDa range. However, important differences exist between the assembly mechanism of these proteins and the mode of action of CL. Multiple PFT families such as hemolysins form large pre-pores at the membrane surface prior to insertion and pore maturation (27). In contrast, CL unrimmed pores present no features above the upper leaflet of the bilayer. Instead, these dynamic structures appear topographically similar to those induced by the antimicrobial peptide melittin and its derivatives (10, 28, 29). The rimmed CL pores are quite distinct from the unrimmed pores. Not only are they more stable on the timescale of AFM imaging (>0.1 s), but rimmed pores have a punctate pattern about their periphery. Such structure is reminiscent of the vestibule of α-hemolysin or the perimeter of pores formed by the N-terminal segment of gasdermin-D (24); however, the overall size of the CL rims is quite modest, only reaching ~0.3 nm outside each side of the bilayer surface. This height corresponds to the difference between the 8-mer (4.6 nm) and the DOPC bilayer thickness (4.0 nm, as measured by AFM, Figure 3 – figure supplement 1). This observation suggests that the pore rims arise when the ends of the TM helices of the 8-mer stick out of the bilayer surface upon 90 degree rotation of the 8-mer in the unrimmed pore, which transitions form the head-to-toe into the side-to-side arrangement (Fig. 8). Some rimmed pores were decorated with protrusions covering the whole pore periphery, forming a corona (Fig. 3B). However, other pores were only partially covered with puncta (Fig. 4A). This observation suggests that maturation into rimmed pores occurs progressively and is not an all-or-none process.

Pore maturation for perfringolysin O (26, 30) and suilysin (20), involves a *vertical collapse* that allows formation of a TM pore. In contrast, the rearrangement of the 8-mer orientation that leads to rim formation causes CL to protrude from the bilayer, and could be defined as a small *vertical swell* between the two types of pores. The 8-mer might rotate because the three K residues located at the Ct (Figure 6A) are expected to be interacting with lipid molecules in the rimmed pore configuration. However, in the vertical 8-mer orientation found in the rimmed pore, the K are expected to be solvent-exposed, resulting in a more stable conformation. An additional difference is that imaging of bilayers in the presence of CL did not show open polymers. However, bilayer removal, either spontaneous or detergent-induced, revealed that linear CL polymers insert into the membranes, but they cannot be readily observed as they do not disturb the bilayer surface or protrude significantly from it. This behavior contrasts with the arcs formed by suilysin, as these rigid open polymers were able to remove lipid molecules from the bilayer to cause perforations (20).

To summarize, here we elucidate how CL forms the membrane pores that cause cellular damage of cells infected by *C. albicans*. Our data show that CL pre-assembles in solution, adopting a pore-competent loop conformation. To the best of our knowledge, this process represents a novel type of molecular mechanism, since in other PFT that form aqueous pores, monomers do not assemble into solution but at the membrane. The assembly of CL into an insertion-competent state in solution might provide an infectivity advantage for *C. albicans*, as it could facilitate faster membrane damage. At the same time, we propose that this assembly mechanism might constitute a therapeutic opportunity to inhibit CL pore formation. Targeting membrane pores presents several drug delivery drawbacks, and it is therefore significantly more challenging than a soluble target. However, drug molecules that prevent CL polymerization in solution could be used as therapeutics to fight *C. albicans* infection. Specifically, a molecule that inhibits the formation of CL loops would be expected to prevent the assembly of the membrane pores that damage human cells.

## Materials and Methods

### Peptide Preparation

Wild-type and variant candidalysin (CL) were synthesized employing solid-phase synthesis by Peptide 2.0 (Chantilly, VA) and HPLC-purified. Purity (>95%) was checked by MALDI-TOF and analytical HPLC. Lyophilized stocks were hydrated in MilliQ H_2_O, stored at −80 °C, and resuspended in the desired buffer at the time of experimentation.

### Liposome Preparation

Lipids were purchased from Avanti Polar Lipids, Alabaster, AL. POPC (1-palmitoyl-2-oleoyl-glycero-3-phosphocholine), DOPC (1,2-dioleoyl-sn-glycero-3-phosphocholine) stocks were prepared in chloroform and stored at −20 °C. Lipids were dried under argon gas and stored in a vacuum overnight prior to use in experiments. Lipid films were resuspended as described below. Large unilamellar vesicles (LUVs) were prepared as described previously (31, 32) using a Mini-Extruder (Avanti Polar Lipids, Alabaster, AL) with a 100 nm filter (Whatman, United Kingdom).

### Fluorescent Dye-Release Assay

Dried lipid films were rehydrated with 50 mM calcein solubilized in 50 mM EDTA and 50 mM NaPi (pH 8) solution. LUVs were formed as described above, separated from free dye by gel filtration using a PD-10 column (GE Life Sciences, Chicago, IL), and diluted to a working lipid concentration of 144 μM. Peptide was added to the calcein entrapped LUVs to achieve the desired lipid:peptide molar ratios. The calcein fluorescence increase caused by dequenching was used as a proxy for CL-induced membrane leakage. A positive control was measured after each treatment, consisting of 0.016% w/v Triton X-100 (TX). A negative control of PBS *in lieu* of peptide treatment was used to calculate percent leakage:

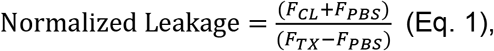

where FI_CL_ is the fluorescence intensity after treatment with CL, FI_TX_ corresponds to the positive control (100% leakage), and FI_PBS_ corresponds to negative control (0% leakage). The quenching effect of TX on calcein was accounted for by measuring the fluorescence intensity of calcein in the presence and absence of TX and calculating a correction factor applied to F_TX_:

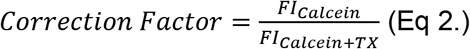

The concentration of calcein was determined by measuring the fluorescence of calcein entrapped within the LUVs post purification. Samples were loaded into a black 96-well plate (Corning, Kennebunk, ME) and measured for two hours, with readings taken every minute, on a Cytation 5 plate reader (BioTek, Winooski, VT) using an excitation wavelength of 495 nm and an emission wavelength of 515 nm.

### Mass Photometry

WT and variant CL were diluted to 333 nM in buffer (10 mM HEPES, 150 mM NaCl, pH 7.3) prior to measurement. Experiments were performed on clean glass coverslips using a Mass Photometer (OneMP, Refeyn). Videos were recorded for 120 s and analyzed on DiscoverMP (Refeyn, version 2.2.0) to determine the molecular weight. The molecular weights were obtained by comparison with known protein standards measured on the same day.

### Analytical Ultracentrifugation

Sedimentation velocity experiments were performed at 20 °C with an An-60Ti rotor in a Beckman Optima XL-I ultracentrifuge at 35,000 RPM. Sedimentation was followed by absorbance at 230 nm and the continuous sedimentation coefficient distribution was obtained using SEDFIT. The following parameters were used for analysis: frictional ratio of 1.2, partial specific volume of 7.3 ml/g, buffer density of 0.9982 g/ml, and buffer viscosity of 1.002 mPa s.

### Atomic Force Microscopy

A sample of one hundred milligrams of DOPC in chloroform was divided into microcentrifuge tubes and dried under argon gas to form uniform films. The tubes were incubated overnight in a vacuum chamber. A dry mechanical roughing pump (XDS5, Edwards) was used to minimize the probability of contamination (11). Samples were back-filled with argon, sealed, and stored at −20 °C. At the time of extrusion, AFM imaging buffer (10 mM Hepes, 150 mM NaCl, pH 7.0) was added to swell the lipids. The lipid solution was extruded (Liposofast, Avestin) through 100 nm membranes 25 times to form unilamellar vesicles. The solution was aliquoted and stored at −80 °C until the time of the experiment. A phosphorus assay was conducted to determine lipid concentration (33). For AFM imaging of CL in solution, stock aliquots of CL were diluted in imaging buffer to the desired concentration (typically 333 nM). A 90 μL droplet was added to freshly cleaved mica and incubated for 10 minutes at 25 °C. Samples were rinsed from above the surface using buffer exchange (90 μL volumes of imaging buffer were exchanged over the surface 5-6 times). Images were collected in imaging buffer with biolever mini tips (Olympus) using tapping mode (Cypher, Asylum Research). Care was taken to keep the magnitude of the tip sample force to ≤100 pN during imaging. As is typical in AFM, lateral image scales are significantly larger than the false color vertical scales.

To image pores, a method for forming lipid bilayers via vesicle rupture was adapted (11). An aliquot of DOPC liposomes was diluted to 300 μM and CL was diluted to specific concentrations The DOPC + CL solutions were incubated in an microcentrifuge tube for 10 minutes, with an additional mixing at the 5 minute mark to encourage peptide diffusion (a pipette was inserted into the solution and the plunger depressed and released 5-6 times). Immediately after incubation in solution, 75 μL of the solution was deposited onto freshly cleaved mica and incubated for another 30 minutes. Material remaining in solution and loosely bound particles were removed by washing via buffer exchange (80 μL volumes of imaging buffer exchanged 5-6 times). All incubations were performed at room temperature (25 °C). Imaging was done in imaging buffer at ~35 °C. The underlying structure of CL peptides was revealed by removing the lipid bilayer with the non-ionic detergent dodecyl beta-D-maltoside (DβM). Stock DβM (196 mM) was diluted in the imaging buffer to 1.5 times the critical micelle concentration (CMC). After imaging pores, the AFM tip was lifted and 75 μL DβM added to the surface and incubated 15 minutes at about 35 °C. To ensure all lipid and detergent molecules were removed from the surface, the samples were heavily rinsed (80 μL volumes of imaging buffer were exchanged 10-12 times). The remaining CL was then imaged in the imaging buffer at 35 °C. Analysis of solution data was performed using commercial software (Asylum Research, Inc.). Pores were analyzed using the Hessian blob algorithm (34). Probability density plots were generated using Epanechnikov kernels and the vertical axes were normalized (integrated to unity area). The number of Gaussian distributions was selected by minimizing the Bayesian information criterion for each model.

For AFM simulations, the tip was modeled as two overlapping spheres of different radii (R = 8 and 4 nm) (11) and the CL 8-mer was positioned in specified orientations. For a single octamer, the geometry of the simulated image (volume = 360 nm^3^) roughly corresponds with the geometry of the smallest subunits observed in AFM (volume of primary experimental peak ±σ= 234 ± 130 nm^3^). Polymeric arrangements of the octamer subunit were made by aligning them head-to-toe in linear or in loop conformations. The inset in **Figure 2C** compares the first four peak locations in the experimental data to the volumes of simulated curved arrangements of 1-4 subunits. Each of the simulated volumes agree with the experimental volume peaks within error.

Curved simulations were chosen for this comparison because a linear model underestimates the measured volume due to a lack of curvature-dependent convolution in the simulation. To avoid volume degeneracies, only small (1-4 subunit) features are analyzed. The persistence length was calculated using Easyworm software (35).

### TEM

Transmission electron microscopy was performed using 2 μM CL samples in AFM imaging buffer incubated on carbon grids for 2 minutes at 25 °C, negatively stained with uranyl acetate, and imaged at 120kV (JEOL, JEM-1400).

### Ion-Mobility Mass Spectrometry

CL peptide samples were directly injected via a syringe pump into a JetStream ESI nebulizer. Mass spectral data and ion mobility measurements were collected using an Agilent 6560 IMS-QTOF mass spectrometer. The peptides were ionized in positive-mode and ions were subsequently pulsed into a helium-filled drift cell in a multi-field fashion (ΔV = 890, 790, 690, 590, and 490 V). The pressure of the drift cell was maintained at 3.940 Torr with the pressure differences between the drift cell and trap funnel being approximately 300 mTorr. An exit funnel and hexapole ion guide focused the ions into the QTOF mass spectrometer. Mobility data were collected over the course of 5.2 minutes (additional instrument parameters are given in **Table S1**). Arrival time distributions were extracted with the Agilent IM-MS Browser software and graphed with Origin Pro.

The ion’s arrival time can be related to the change in drift cell voltage (ΔV) to determine the ion’s reduced mobility (*K*_0_). To calculate the momentum transfer collision integral, which approximates the experimental collisional cross section (*σ_exp_*), the Mason-Schamp equation was used (36):

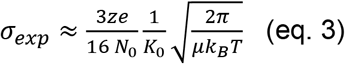

Where z is the ionic charge, e is the elementary charge, *N_0_* is the number gas density, μ is the reduced mass of the ion-buffer gas pair, *k_B_* is the Boltzman constant, and T is the temperature of the buffer gas. Theoretical collisional cross sections were calculated using the trajectory method available in the Mobcal package (37).

### Molecular Dynamics Simulation

The atomistic model of CL 8-mer was built by aligning the peptide sequence to that of the eight-helix coiled coil CC-Type2-II (PDB ID 6G67) (38) using EMBOSS (39). The initial structure was minimized using the GROMACS 4.6.7 (40) during 25 ns and the CHARMM27 force field. Molecular dynamics (MD) simulation of the dimer was performed using the same package and force field. The initial structure was minimized using the steepest decent method and solvated in a TIP3P cubic water box (a = 14.03 nm). Chloride anions were added to neutralize the charges. Solvent and volume equilibration simulations in NPT ensemble (T = 300K and P = 1bar) were performed to optimize the box size, followed by 45-ns NPT simulations at 300K. The LINCS algorithm (41) was employed to constrain bonds between heavy atoms and hydrogen, and the SETTLE algorithm (42) was used for water molecules. These constraints allowed an integration time step of 2.0 fs. The electrostatic and dispersion forces were computed with a real space cutoff of 1.2 nm, and particle mesh Ewald method (43) was used to treat long range electrostatics. The temperature was maintained by the Nose-Hoover thermostat. The temperature and pressure coupling constants were 0.1 and 1.0 ps, respectively. The equations of motion were integrated according to the leap-frog algorithm.

### Circular Dichroism Spectroscopy

The circular dichroism (CD) spectrum of candidalysin was collected in the presence and absence of liposomes. LUVs were prepared by extruding resuspended lipid at 1 mM in sodium phosphate buffer pH 7.4. The final lipid concentration was 130 μM, and the final peptide concentration was 5 μM. CD spectra were measured in a Jasco J-850 spectropolarimeter with a 2 mm path length. A lipid blank and a buffer blank were subtracted from the lipid and lipid-free samples. Mean residue ellipticity (MRE) was calculated as:

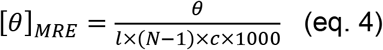

where *θ* is ellipticity (millidegrees), *l* is cuvette pathlength (mm), *N* is the number of amino acids in the peptide, and *c* is the concentration of the peptide (M).

### Oriented Circular Dichroism

Two circular quartz slides (Hellma Analytics, Müllheim, Germany) were cleaned by submerging in piranha solution (75% H_2_SO_4_, 25% H_2_O_2_) for five minutes. POPC/POPE/cholesterol films were resuspended in 2,2,2-trifluoroethanol with or without CL, and this mixture was dried onto the slides for at least twelve hours at room temperature and pressure. The amount of lipid on each slide was 85.3 nmol, and the amount of peptide on each slide 1.71 nmol, for a lipid to peptide ratio of 50:1. The deposited lipid was then rehydrated with PBS in a chamber containing saturated potassium sulfate, which produces a relative humidity of 96% to prevent the films from dehydrating. After at least twelve hours, the slides were loaded onto an OCD cell containing saturated potassium sulfate. OCD spectra were measured eight times on a Jasco J-815 spectropolarimeter at RT. The OCD cell was rotated 45° between measurements to correct for any imperfections in the lipid bilayers. To obtain the final spectra, the eight measurements were averaged, and the peptide-free lipid blanks were subtracted.

### Cell Culture

Experiments were performed using the TR146 buccal squamous carcinoma cell line obtained from the European Collection of Authenticated Cell Cultures (ECACC 10032305) and grown in Dulbecco’s Modified Eagle Medium (DMEM, Gibco) supplemented with 10% fetal bovine serum (FBS) and 1% penicillin-streptomycin. All experiments were performed in serum-free DMEM.

### Western Blot

TR146 cells were grown to confluency and starved overnight in serum-free DMEM. Cells were treated with 15 μM peptide for two hours. Post-incubation, cells were lysed on ice with TEN-T buffer (50 mM Tris-HCl pH 7.5, 100 mM NaCl, 1 mM EDTA, 1% Triton-X 100) containing phosphatase (Sigma) and protease (ThermoFisher) inhibitors for 30 minutes. Lysates were collected post centrifugation (20 minutes, 13,000 RPM, 4 °C) and separated on 10% SDS-PAGE gels before transfer to 0.45 μm nitrocellulose membranes. Blots were probed with primary antibodies for c-Fos (1:500, Cell Signaling Technology, 2250S) and phospho-MKP1 (1:1000, Cell Signaling Technology, 2857S) overnight. Fluorescent secondary antibodies (1:10,000, goat-anti mouse IRDye 680, goat-anti rabbit IRDye 800, LI-COR) were used for detection on a LI-COR Odyssey CLx. Human ß-actin (1:1000, Abcam, ab6276) was used as a loading control and protein expression and phosphorylation was quantified using ImageStudio software.

## Acknowledgements

We thank Jennifer Schuster (University of Tennessee) for comments on the manuscript, and Ed Wright (University of Tennessee) for assistance with the AUC experiments. We are also thankful to Erwin London for scientific advice. This work was partially funded by awards NIH R35GM140846 and R01GM120642 (to F.N.B) and NSF 1709792 and 2122027 (to G.M.K.).

## Notes

### Competing Interest Statement

The authors have declared no competing interest.

