## Supplementary Material for "The *C. albicans* virulence factor Candidalysin polymerizes in solution to form membrane pores and damage epithelial cells"

### Supplementary Materials

10 figures
1 table

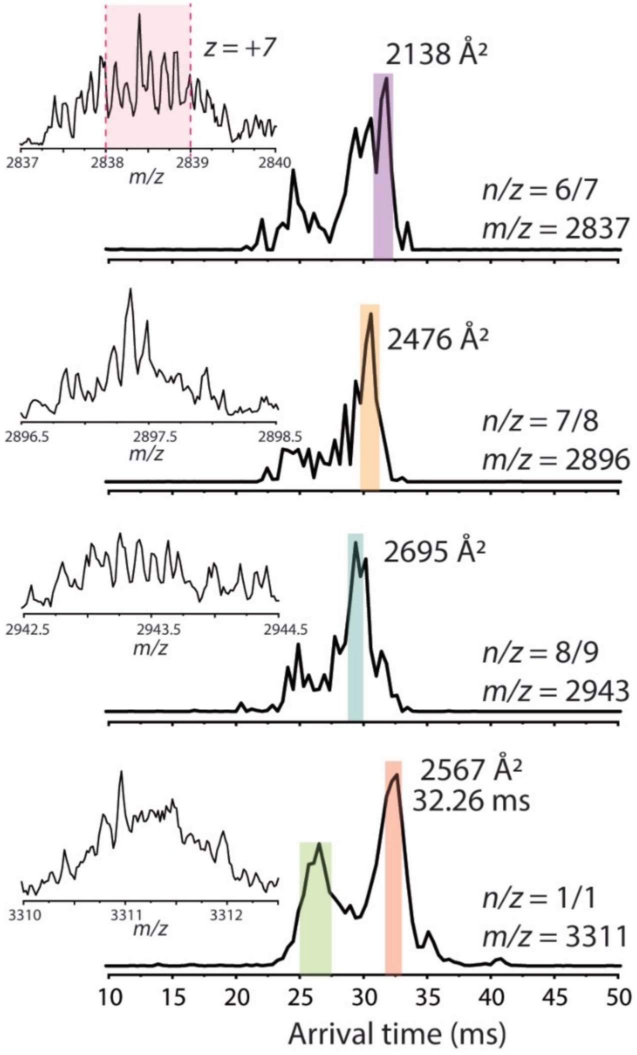


**Figure 1 – figure supplement 1.** Arrival time distribution of higher-ordered Candidalysin oligomers. The experimental collisional cross sections for prominent features (highlighted regions) are given. The inset above each ATD details the mass spectrum of each respective species with calculated isotopic spacing. The highlighted region within the mass spectrum of *n/z* = *6/7* emphasizes that this species unambiguously has a charge state of +7 due to isotopic spacing. The last ATD shows a species with a nominal oligomer-to-charge ratio of *1/1*. However, by comparing the isotopic spacing(s) with other mass spectral peaks, the species at *m/z* 3311 must be *at least* an octamer with *z* = +*8* (i.e. *n/z* = 8/8) since anything less than *z* = 8 should be resolved. We conservatively assigned the longer arrival time feature (red highlight) to an octamer (*n/z* = 8/8). To further support this assignment, we note that the species at *n/z* = 7/8 and 8/9 exhibit faster arrival times compared to *n/z* = 8/8. This follows the expectant logic since *n/z* = 8/9 has a higher charge state than *n/z* = 8/8, and thus traverses the drift cell faster. On the other hand, *n/z* = 7/8 has the same charge state as *n/z* = 8/8, but is a smaller oligomer, which results in a shorter arrival time.

| | Parameter | Value |
|---|---|---|
| Pre-IMS Zone | Source: Gas Temperature | 300 $^{o}$C |
| | Source: Drying Gas | 5 L/min |
| | Source: Nebulizer Pressure | 13 psi |
| | Source: Capillary | 3800 V |
| | Optics I: Fragmentor | 250 V |
| | IM Front Funnel: High Pressure Funnel Delta | 110 V |
| | IM Front Funnel: High Pressure RF Delta | 180 V |
| | IM Front Funnel: Trap Funnel Delta | 160 V |
| | IM Front Funnel: Trap Funnel RF | 180 V |
| | IM Front Funnel: Trap Funnel Exit | 10 V |
| | IM Trap: Trap Entrance Grid Low | 82 V |
| | IM Trap: Trap Entrance Grid Delta | 2 V |
| | IM Trap: Trap Entrance | 79 V |
| | IM Trap: Trap Exit | 76 V |
| | IM Trap: Trap Exit Grid 1 Low | 72 V |
| | IM Trap: Trap Exit Grid 1 Delta | 6 V |
| | IM Trap: Trap Exit Grid 2 Low | 71 V |
| | IM Trap: Trap Exit Grid 2 Delta | 13 V |
| | Acquisition: Trap Fill Time | 1000 µs |
| | Acquisition: Trap Release Time | 100 µs |
| Post-IMS Zone | IM Drift Tube: Drift Tube Exit | 210 V |
| | IM Rear Funnel: Rear Funnel Entrance | 200 V |
| | IM Rear Funnel: Rear Funnel RF | 130 V |
| | IM Rear Funnel: Rear Funnel Exit | 35 V |
| | IM Rear Funnel: IM Hex Entrance | 42 V |
| | IM Rear Funnel: IM Hex Delta | -8 |
| | Optics 1: Oct Entrance Lens | 32 V |
| | Optics 1: Lens 1 | 28.3 V |
| | Optics 1: Lens 2 | 15.8 V |
| | Quad: Quad DC | 26.6 V |
| | Quad: Postfilter DC | 26.5 V |
| | Cell: Gas Flow | 22 psi |
| | Cell: Cell Entrance | 25.6 V |
| | Cell: Hex DC | 24.2 V |
| | Cell: Hex Delta | -9 V |
| | Cell: Hex2 DC | 15 V |
| | Cell: Hex2 DV | -3 V |
| | Optics 2: Hex3 DC | 11.8 V |
| | Extractor: Ion Focus | 5.6 V |

**Table Supplement 1. Agilent 6560 IMS-QTOF Parameters**

**A**

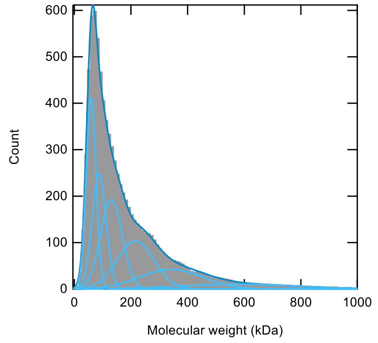


**B**

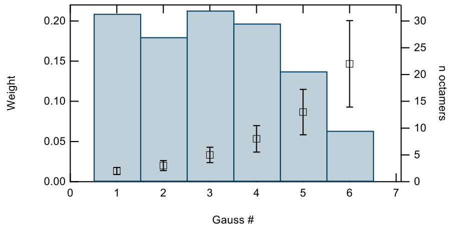


Mass, kDa

| | **Gauss 1** | **Gauss 2** | **Gauss 3** | **Gauss 4** | **Gauss 5** | **Gauss 6** |
|---|---|---|---|---|---|---|
| **Mass, kDa** | **59.05** | **87.07** | **130.50** | **215.32** | **341.32** | **578.13** |
| **Amplitude** | 411.77 | 248.90 | 191.39 | 103.23 | 42.32 | 10.51 |
| **HWHM** | 20.60 | 29.32 | 45.17 | 77.39 | 131.87 | 251.85 |
| **Std. Dev.** | 17.50 | 24.91 | 38.36 | 65.72 | 112.00 | 213.90 |
| **8-mer N** | **2** | **3** | **5** | **8** | **13** | **22** |

**Figure 1– figure supplement 2. (A)** Gaussian fitting of the MP data of CL WT. The table shows the fitting parameters and the resulting errors. BIC (Bayesian information criterion) analysis was used to determine the optimal number of Gaussian terms for the fitting. **(B)** The relative weight of the six Gaussian components shows that the first peaks, corresponding to two, three, five and eight 8-mers, are the most abundant species, and are similarly populated.

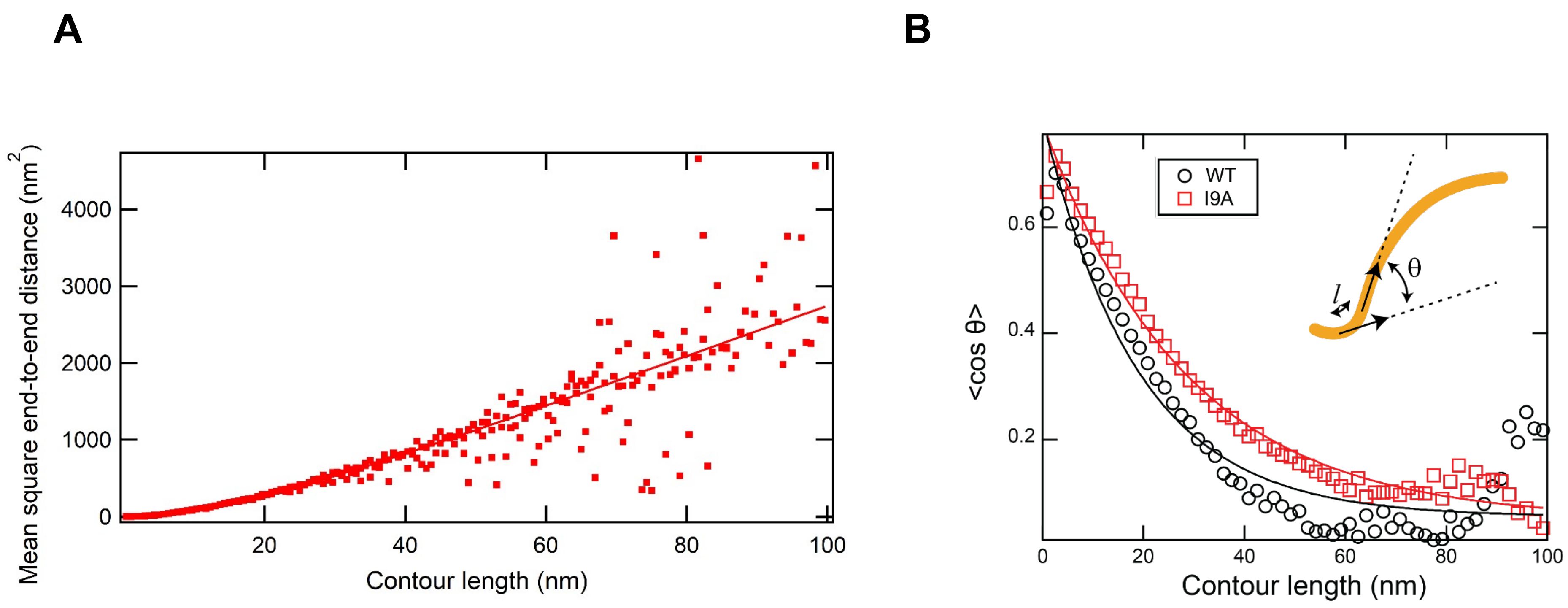


**Figure 2 – figure supplement 1. (A)** AFM images of CL in solution yield a persistence length, $L_p$ , of 9 ± 2 nm. The value was calculated from a fit (solid line, coefficient of determination = 0.87) to the mean square end-to-end distance of *N* = 100 polymers. **(B)** An identical value of $L_p$ (within uncertainty of the analysis) was obtained by calculating the decay of tangent-tangent correlations. I9A in solution yielded a persistence length, $L_p$, of 13 ± 2 nm. The value was calculated from the exponential decay (*solid line*) of tangent-tangent correlations of *N* = 100 polymers. *Inset*: The angle $\theta$ is defined by polymer segments separated by a distance *l* along the contour of the polypeptide

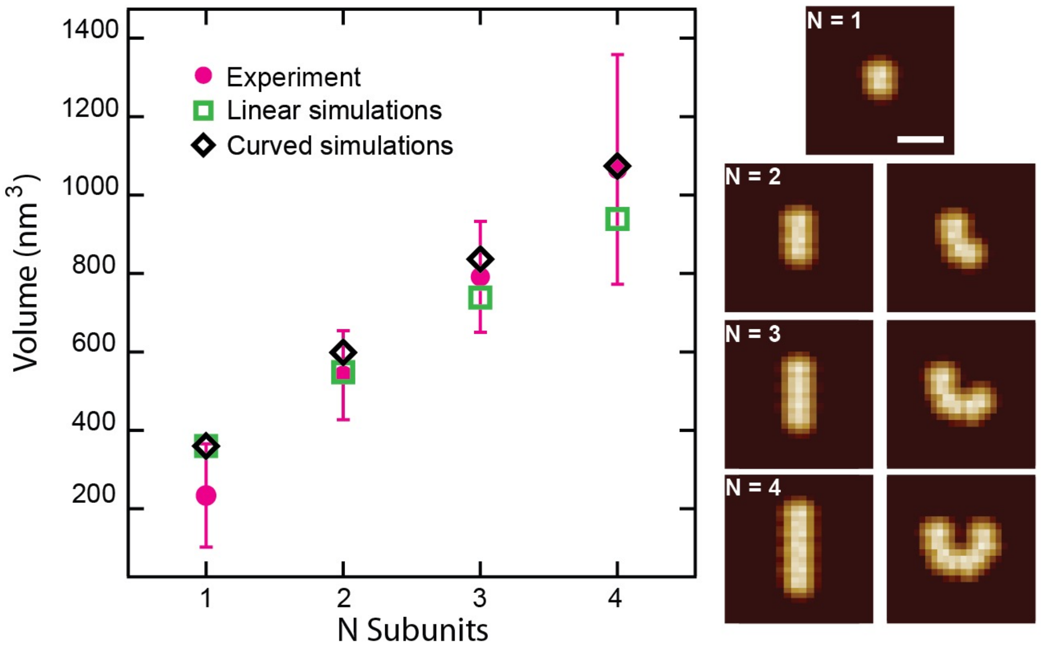


**Figure 2 – figure supplement 2.** Comparison of N 8-mer subunits in the side orientation arranged either linearly or with an added curvature (Scale bar = 15 nm). Volumes are plotted for each particle showing larger volumes for curved polymers. Error bars for the peak positions represent the standard deviation.

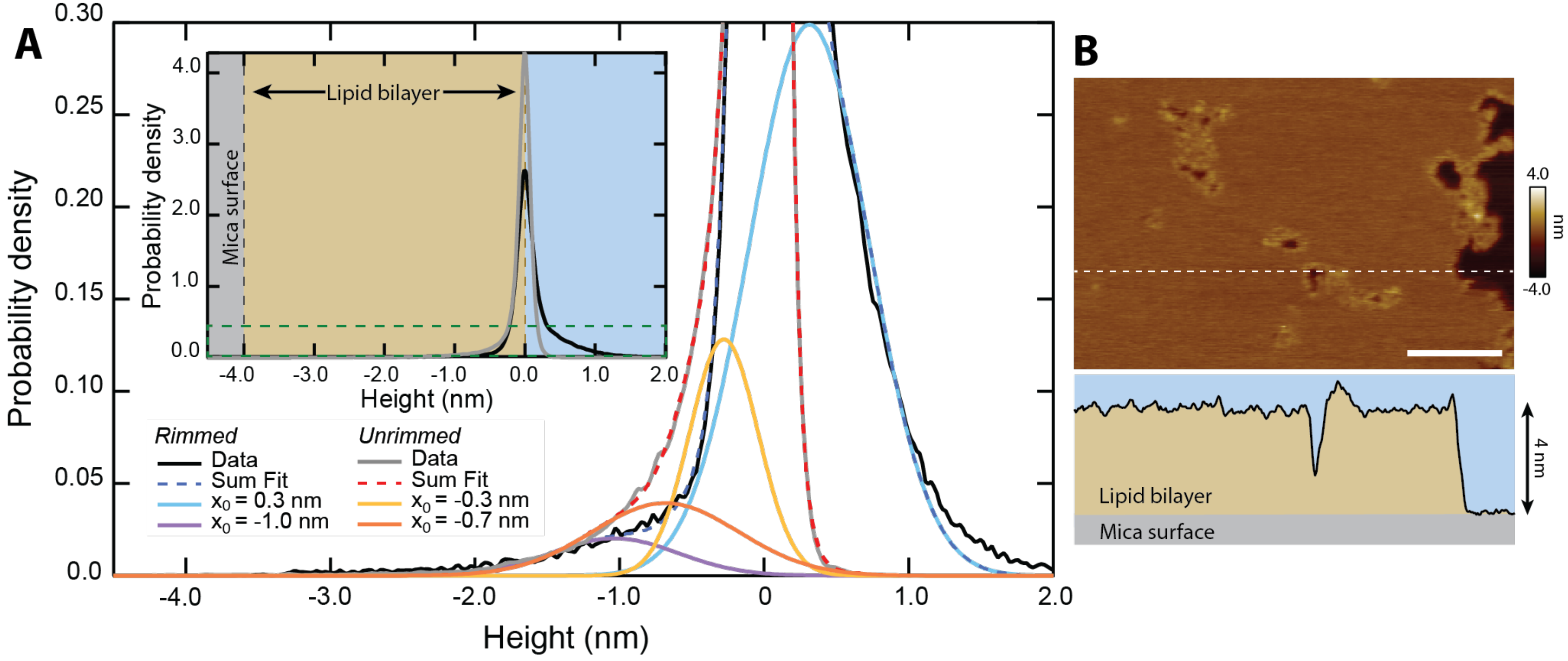


**Figure 3 – figure supplement 1. (A)** The inset shows the height distribution of all pixels for *N* = 313 unrimmed pores (gray, *n* = 153,832 pixels) and *N* = 193 rimmed pores (black, *n* = 114,996 pixels). Zero height represents the top of the upper leaflet of the lipid bilayer. The main graph is rescaled to show the area encompassed by the dark green dashed line. Multiple Gaussians are fitted to the histograms to deconvolve the different populations. Both unrimmed and rimmed pores had populations that deviated from the background (~0 nm), which are plotted in addition to the summary fits. The unrimmed pores show two distinct depths, -0.7 ± 0.5 nm and -0.3 ± 0.2 nm (mean ± σ.) The rimmed pores have a single population lower than the background, located at -1.0 ± 0.4 nm. Additionally, a positive population is located at +0.3 ± 0.4 nm, corresponding to the topographically high rims. **(B)** A line scan (white dashed line) across an image of rimmed pores demonstrates the 4 nm depth of the DOPC bilayer in profile; scale bar = 100 nm.

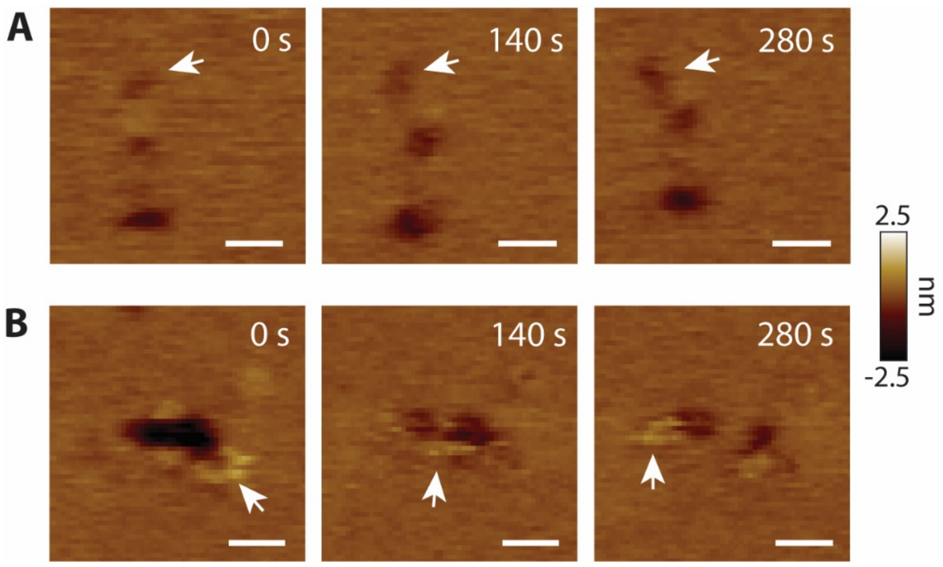


**Figure 3 – figure supplement 2. CL pores exhibit lateral dynamics in the AFM supported membrane (A)** Lateral diffusion of an unrimmed pore (arrows). **(B)** A void forms two pores. The arrows highlight positive features; scale bars = 20 nm.

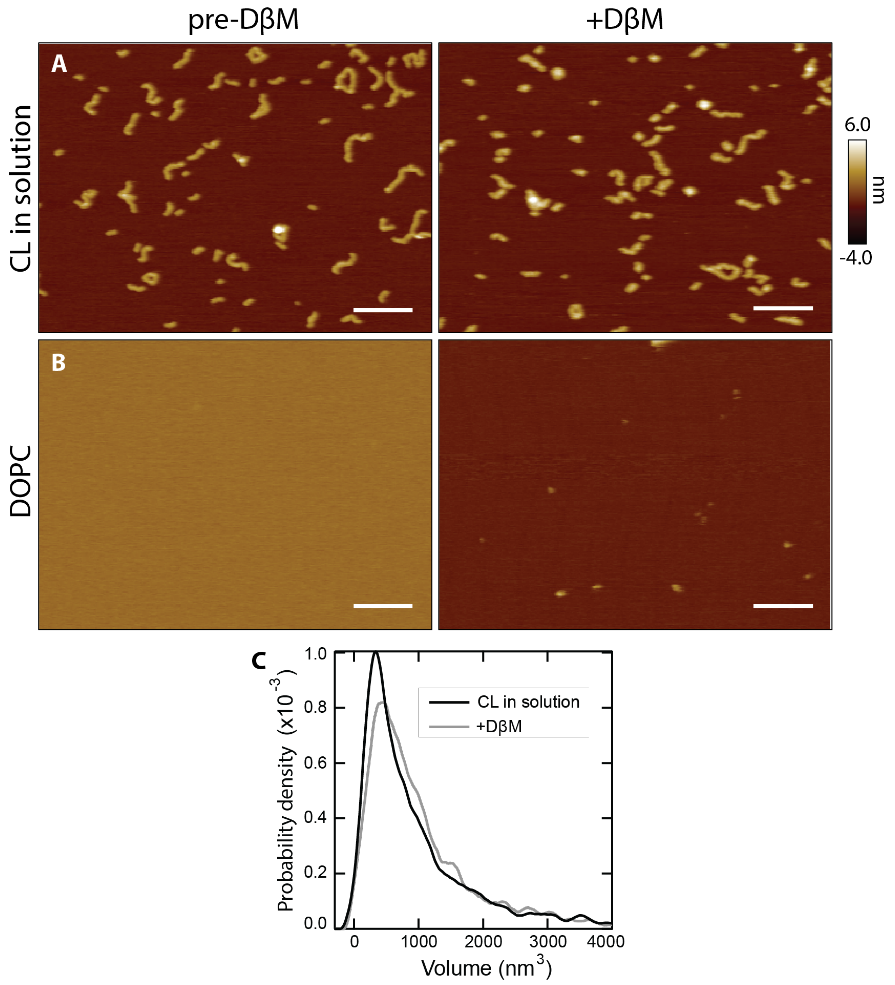


**Figure 4– figure supplement 1.** (**A**) Adding DβM to CL in solution (i.e., in the absence of lipid) induces no significant change in CL polymers, which remain adhered to the mica surface (scale bars = 100 nm). **(B)** DβM added to bare DOPC bilayers removes the bilayer with a small amount of residual lipid left behind on the mica surface. **(C)** A volume histogram of CL after DβM addition (gray) shows a slight increase in size of the features compared to the CL in solution data (black), likely attributable to occasional detergent molecules binding to CL.

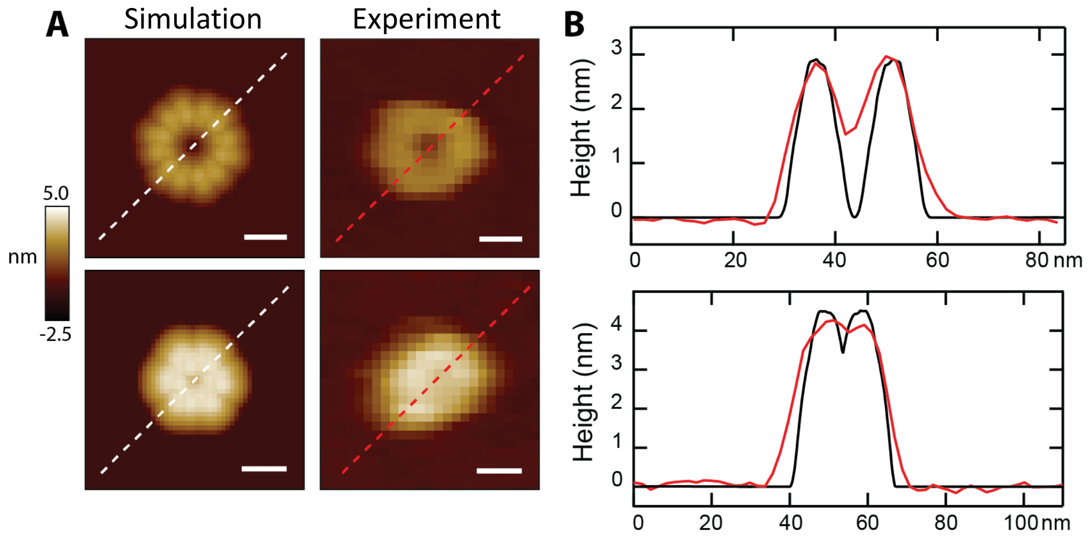


**Figure 5, figure supplement 1. Area analysis and simulation studies. (A)** Simulated images of loops formed by six 8-mers are shown side-by-side with experimental data compatible for the two orientations. The upper panel compares a head-to-toe simulation with a solution loop:, and the lower panel shows the vertical model compared to a CL pore after lipid  lipid spontaneously dissociated. **(B)** Line scans through the feature demonstrate geometric agreement between models (black lines) and experimental data (red lines).

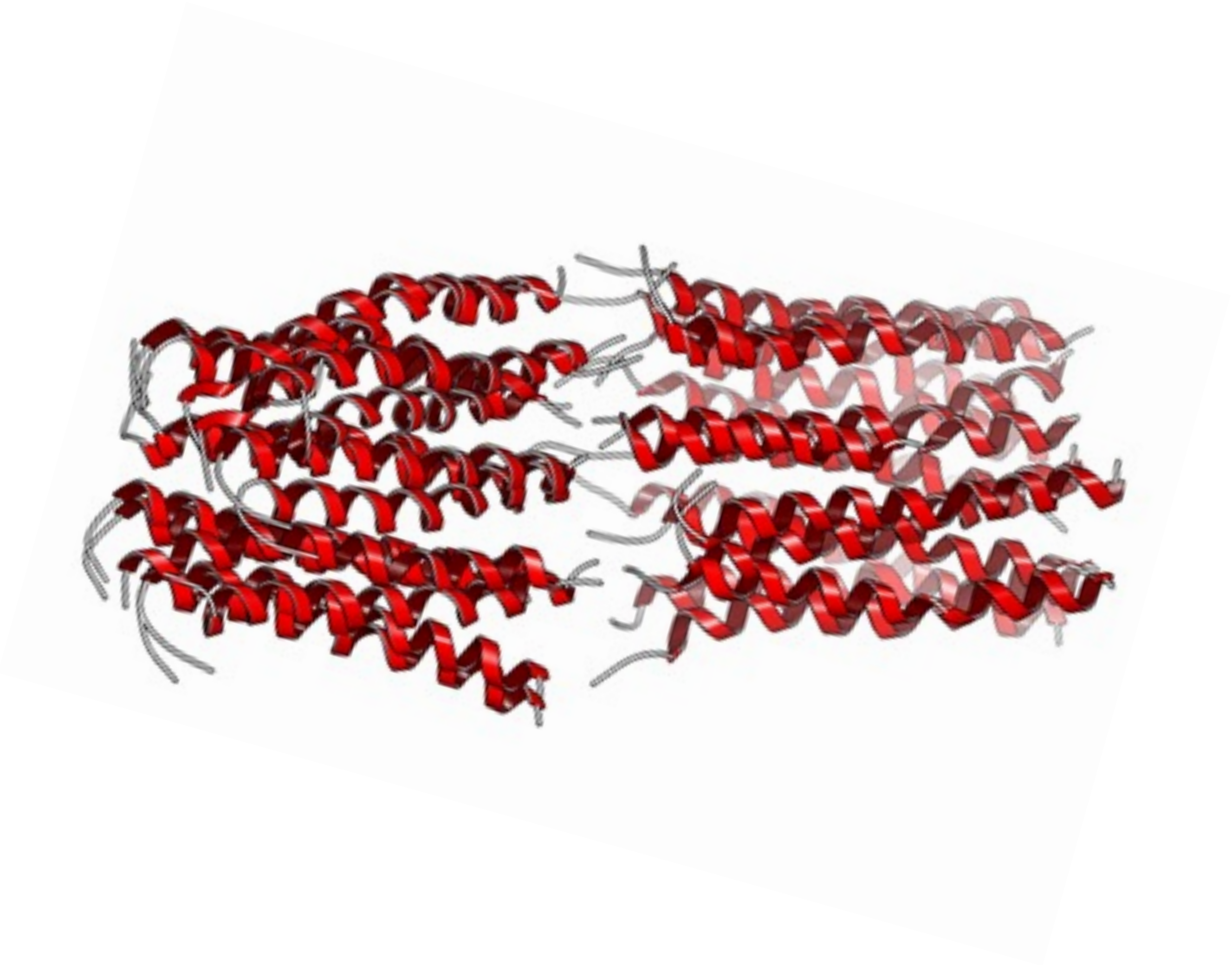

**Figure 6– figure supplement 1. Modeling of the assembly between two CL 8-mers.**

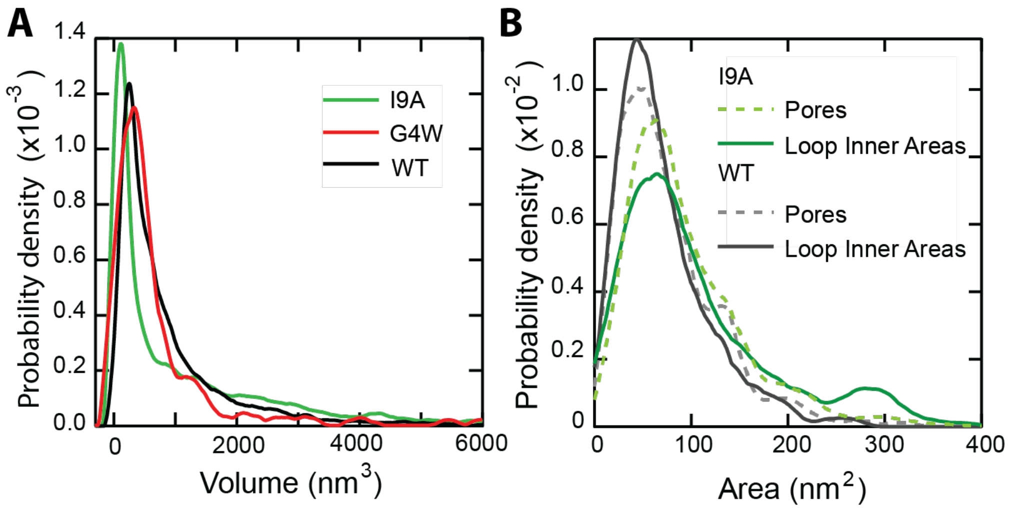


**Figure 6– figure supplement 2. (A)** A volume histogram of CL in solution compares the wildtype ($N$ = 7,838 features), I9A ($N$ = 3,609), and G4W ($N$ = 422). The I9A mutant has a smaller primary peak, ~100 nm$^3$, indicating a large number of smaller features. The larger volumes of the loops fall into the long high-volume shoulder. **(B)** An area histogram of $N$ = 1397 unrimmed pore-like features shows I9A increases the pore area of the primary peak population >20%, from 50 nm$^2$ (unrimmed wild type, $N$ = 1468) to 61 nm$^2$. $N$ = 156 I9A loops are analyzed with the Hessian blob algorithm to get the inner areas and compared to the unrimmed I9A pore areas ($N$ = 1397). The overall agreement between pore geometry and loop inner areas for both the I9A mutant and the WT ($N_{rings}$ = 261) CL indicate that structures formed in solution can directly translate into membrane pores.

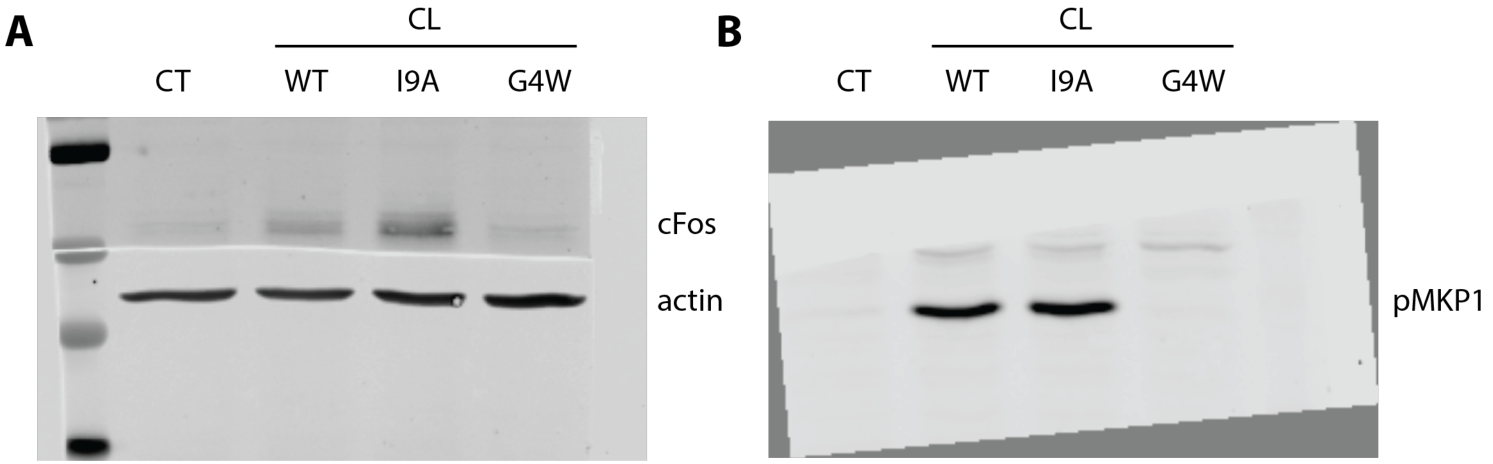


**Figure 7– figure supplement 1.** Uncropped Western blot of TR146 cell lysates treated with WT and variant CL for two hours probed with antibodies for (A) cFos, actin and (B) pMKP1.
